# Modelling geospatial distributions of the triatomine vectors of *Trypanosoma cruzi* in Latin America

**DOI:** 10.1101/738310

**Authors:** Andreas Bender, Andre Python, Steve W. Lindsay, Nick Golding, Catherine L Moyes

**Author notes:** Big Data Institute, Li Ka Shing Centre for Health Information and Discovery, University of Oxford, Old Road Campus, Oxford, UK.

## Abstract

Approximately 150 triatomine species are known to be infected with the Chagas parasite, *Trypanosoma cruzi*, but they differ in the risk they pose to human populations. The largest risk comes from species that have a domestic life cycle and these species have been targeted by indoor residual spraying campaigns, which have been successful in many locations. It is now important to consider residual transmission that may be linked to persistent populations of dominant vectors, or to secondary or minor vectors. The aim of this project was to define the geographical distributions of the community of triatomine species in Latin America. Presence-only data with over 12, 000 observations of triatomine vectors were extracted from a public database and target-group background data were generated to account for sampling bias in the presence data. Geostatistical regression was then applied to estimate species distributions and fine-scale distribution maps were generated for thirty triatomine vector species. The results for *Panstrongylus geniculatus, P. megistus, Triatoma barberi, T. brasiliensis*, and *T. pseudomaculata* are presented in detail and the model validation results for each of the 30 species are presented in full. The predictive maps for all species are made publicly available so that they can be used to assess the communities of vectors present within different regions of the endemic zone. The maps are presented alongside key indicators for the capacity of each species to transmit *T. cruzi* to humans. These indicators include infection prevalence, evidence for human blood meals, and colonisation or invasion of homes. A summary of these indicators shows that the majority of the 30 species mapped by this study have the potential to transmit *T. cruzi* to humans.

**Author summary:** The Pan American Health Organisation’s Strategy and Plan of Action for Chagas Disease Prevention, Control and Care highlights the importance of eliminating those triatomine vector species that colonise homes, and has had great success in many locations. Since indoor residual spraying campaigns have targeted these species, their importance relative to other vectors has diminished and their geographical distributions may also have changed. It is now vital to consider the full community of vector species, including previously dominant vectors as well as secondary or minor vector species, in order to target residual transmission to humans. Our aim was to define the geographical distributions of the most commonly reported triatomine species in Latin America. We extracted reports of triatomine vector species observed at specific locations from a public database and we used a geostatistical model to generate fine-scale predictive maps for thirty triatomine vector species. We present these maps alongside a summary of key indicators related to the capacity of each species to transmit the Chagas parasite to humans. We show that most of the 30 species that we have mapped pose a potential threat to human populations.

## Introduction

American trypanosomiasis, or Chagas disease, is one of the 10 neglected diseases addressed by the London Declaration, which calls for control and elimination of these devastating diseases by 2020 [1]. It is a disease of Latin America and the Pan American Health Organisation (PAHO) has set out a ‘Strategy and Plan of Action for Chagas Disease Prevention, Control and Care’ [2]. This strategy includes the elimination of domestic vectors to prevent intra-domiciliary transmission, as well as screening blood donors and pregnant women to prevent transmission via blood donation or the placenta, and implementation of best practice in food handling to prevent oral transmission. Our study focuses on the primary route of infection; the contamination of a vector bite by the faeces of that vector.

The *T. cruzi* parasite is transmitted to humans by over 150 different vector species from 18 different genera [3]. The transmission risk that each vector species poses is influenced by how likely it is to come into contact with humans and this, in turn, is influenced by short-distance movement (for example does the species enter and/or colonise homes) and its larger-scale geographical distribution. Studies assessing vulnerability of individuals to Chagas disease have shown that, while housing, ecotype and soci-economics are all relevant, triatomine presence is the most important indicator [4]. Thus understanding the distribution of these vector species is vital to both target control measures and to assess disease risk.

Before the current intervention era, five vector species were recognised as being dominant in the transmission of *Trypanosoma cruzi* to humans based on their habit of colonising houses, behaviour (feeding-defecation interval) and widespread geographical distributions [5]. Since indoor residual spraying (IRS) campaigns have successfully targeted these dominant species in many locations, their importance relative to other vectors has diminished and their geographical distributions may also have changed [6]. It is now vital to understand the full community of vector species, including previously dominant vectors as well as secondary or minor vector species, in order to target residual transmission to humans [6–8]. Data on intervention coverage with spatially and temporally high resolution across the entire zone was, however, unavailable and thus could not be taken into account at the modelling stage.

Several studies have investigated species behaviours that influence short distance travel in and around homes, such as host-seeking, aggregation and dispersal [9–19], but fewer studies have considered the larger-scale geographical distributions of these species. The studies of geographical species distributions that have been conducted typically focus on a single country or a region within a country [19–24]. One earlier study considered the distribution of infected vectors across South America, without distinguishing species [25], but no studies have considered the geographical distributions of individual dominant and secondary vector species in Latin America. A lack of consistent region-wide information makes it harder to construct an overview for the region as a whole or to compare areas within the endemic zone.

The data recording presence of a species are often sparse and suffer from sampling bias, which makes inter-region comparison of these records difficult. The aim of this study is to use statistical models to produce a comprehensive set of maps predicting the distributions of triatomine vector species while taking into account the limitations of the data. We use an extensive database of reported occurrences of each species, data on environmental variables that are likely to influence species presence, and build species distribution models to improve our current understanding of the spatial distribution of *T. cruzi*.

## Materials and methods

### Species occurrence and background points

The primary source of vector species data was a database of vector occurrence locations, which was supplemented with additional species presence points derived from a database of infections in vector species. Data on vector occurrence were extracted from DataTri, a publicly available database that reports the presence of a given triatomine species, the date of collection (if available) and geographical coordinates for each collection [26]. Additional vector occurrence data was added using a database of *Trypanosoma cruzi* infections in triatomines that also provided the vector species found, the date of collection (if available) and geographical coordinates for each collection [27]. Any data points from DataTri that were duplicated in the second data set were removed before vector occurrence data from the infections database were added to the DataTri data set. Data points before the year 2000 were removed because the aim was to investigate vector distributions in the current era.

The available vector occurrence data is usually referred to as *presence only* data. Techniques for modelling such data often involve augmenting the presence data with *pseudo-absence* or *background* points, which requires a source of appropriate background data [28–30]. Here we use a target-group background (TGB) approach by choosing background data that exhibits similar sampling bias as the occurrence data [31]. This approach can reduce the bias introduced by preferential sampling of the presence locations. It was successfully used to map geographical distributions of malaria hosts and vectors [32] and predict infection risk zones of yellow fever [33]. In simulation studies this method also performed well when compared to approaches using presence-absence data [31, 34]. As with all models of presence-only data, the maps produced using the TGB approach represent relative rather than absolute probabilities of species occurrence.

We constructed one TGB data set for each vector species as outlined below and illustrated in Figure 1 for *Panstrongylus megistus*:

**Fig 1.**
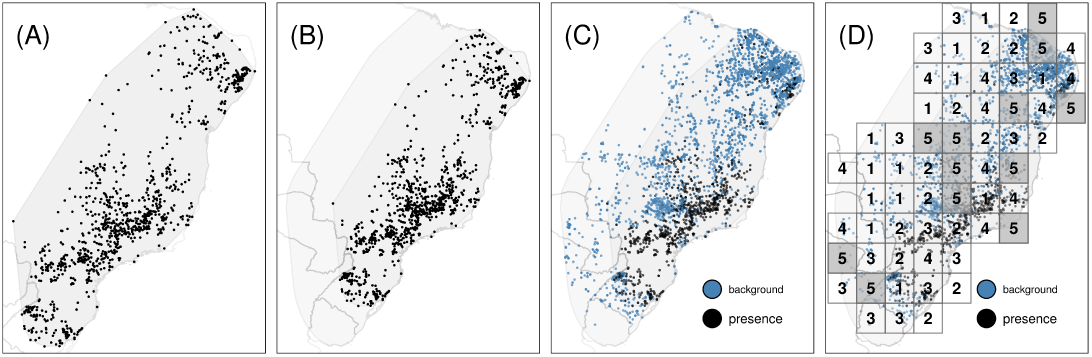
Construction of background points. Illustration of the construction of background points using the TGB approach for species *Panstrongylus megistus*. Panel (A): A convex hull is constructed around the presence locations of the species. Panel (B): The hull is extended by a fixed width of 5 degrees (*extended hull*). Panel (C): Background points are added using presence locations of all other species within the extended hull. Panel (D): Blocks of width *w*_*k*_ are allocated randomly across the extended hull. The observations in blocks numbered 1-4 are used as training data, and the observations in blocks numbered 5 are assigned to the test data. One fold consists of all blocks sharing the same number.

1. The presence locations of vector species *k* = 1, …, *K* (target-group) were extracted from the database and a convex hull containing all presence locations was constructed (panel (A) in Figure 1).
2. This hull was extended by a constant width of 5 degrees in all directions to allow for uncertainty with respect to the range of the species being modelled (*extended hull*; panel (B) in Figure 1).
3. The presence locations of all other species within the extended hull were defined as background points (blue dots in panel (C) of Figure 1).
4. Duplicate observations at the same site and the same year were removed.
5. At the modelling stage, the background points were weighted such that their total weight is equal to the number of presence observations (cf. [31, 32]).

For some species, only few observations were available in the data set. It was suggested that approximately five [35] or ten [36–38] events (presences) per predictor are required to reliably fit a logistic regression. Given that we use up to 30 predictors, this would imply a sample size of *n* ≈ 150 and ≈ 300, respectively. In our data, 14 and and 9 species fulfilled this (approximate) requirement. For completeness, we fit models for all species with *n* > 50, but obviously results must be interpreted with care as the sample size (number of presence observations) decreases (see Results section). Species with fewer than fifty observations in the training and test data were not modelled, however, their presence locations were used as background points for the species that were modelled.

### Environmental variables

Previous work has shown that vector distributions are influenced by climate, land cover types and rural/urban classifications [19–24]. Environmental variables for these three data types were obtained at a nominal resolution of 5 × 5 kilometres. The climatic variables used were *land surface temperature* (annual; day, night and diurnal difference) [39], two measures of *surface moisture* (annual) [40], *rainfall* (annual) [41], *elevation* (static) [42] and *slope* (static) [43]. The variable used for land cover were the 16 IGBP *land cover* classes (annual) [44] and an *enhanced vegetation index* [45]. *Finally the variables used to distinguish rural, peri-urban and urban areas were urban footprint* (static) [46], *nighttimelights* (static) [47], *human population* (annual) [48] and *accessibility* (static, based on road networks and distance to cities) [49]). Annual environmental variables were not always available for all time periods in which occurrence data was available. In this case, we instead used values from the closest year.

### Model evaluation

In the context of spatial analysis, data available for modelling often only encompasses few locations in areas for which predictions are generated. Therefore, the model is usually evaluated on out of sample data to avoid over-fitting, to ensure transferability to new locations and to obtain realistic estimates for the goodness of fit. Standard approaches to model evaluation, however, can yield over-optimistic metrics of the model predictive ability unless the spatial nature of the data (and the model) is taken into account [50–52]. To address these concerns, the data was initially split randomly into *train-test data* (80%) and *evaluation data* (20%), stratified by species. The latter data set is only used for the final model evaluation, without ever being used for estimation (column “AUC*” in Table 2). Additionally, the train-test data was split into five folds. Following recommendations in [53] each fold consisted of multiple spatial blocks, where the block size *w*_*k*_ for species *k* = 1, …, *K* was set such that approximately 50 blocks (10 per fold) would cover the extended hull of that species and defined as 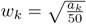, where *a*_*k*_ is the area of the extended hull of species *k* = 1, …, *K*. Figure 1 (panel (D)) depicts the resulting blocks and folds for species *Panstrongylus megistus*. For each species, blocks one through four were assigned to the training data, while blocks numbered five (grey shade) were only used to obtain out-of-sample test errors. Allocation of blocks was spatially random to avoid systematic bias of presence and background locations in any of the folds, but stratified with respect to species presence such that the proportion of presence and background points was approximately equal in all folds. The spatial blocking for all species considered in our analyses are provided in the supplement (S1 File). Model performance was evaluated by the area under the receiver operator curve (AUC), which measures the models ability to discriminate between presence and background points.

### Modelling

Let *ε*_*k*_ ∈ ℝ^2^ the extended hull of species *k* = 1, …, *K* and *y*_*k,t,i*_ ∈ {0, 1} ∼ *Bernoulli*(*π*_*k,t,i*_), *i* = 1, …, *n*_*k*_ the binary indicator of presence/background for species *k* in year *t* ∈ *{*2000, …, 2016} at location *s*_*i*_ ∈ *ε*_*k*_ (as constructed by the TGB approach). The relative probability of occurrence *π*_*k,t,i*_ is estimated by a logistic generalised additive model (GAM) with linear predictor (1)

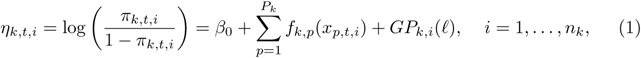

where *f*_*k,p*_(*x*_*p,t,i*_) is the species specific, potentially non-linear, effect of the *p*-th covariate estimated by a penalised thin-plate spline [54] and *GP*_*k,i*_(ℓ) is a two-dimensional, species specific Gaussian process (GP) with range parameter ℓ evaluated at location *s*_*i*_ (the smoothness parameter was set to 1.5). The number of covariates *P*_*k*_ can vary by species as some of them might not have enough unique values within the spatial extent of the species to be relevant for analysis. Here, covariates were only included if the number of unique values was at least twenty. The correlation function of the GP was defined by *C*(*x, x*′) = *ρ*(‖*x - x*′‖), where *ρ*(*d*) = (1 + *d/*ℓ) exp(*-d/*ℓ) is the simplified Matérn correlation function with range parameter ℓ = *max*_*ij*_ ‖*x*_*i*_ − *x*_*j*_‖ as suggested in [55] and implemented in [56].

The model was estimated by optimising the penalised restricted maximum likelihood (REML) criterion (2)

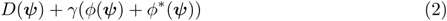

using a double shrinkage approach where ***ψ*** is a vector of all coefficients associated with the smooth functions *f* and *GP*, *D*(***ψ***) is the model deviance and *ϕ*(.) and *ϕ*\*(.) are range space and null space penalties of the model coefficients ***ψ*** [54, 57]. The first penalty (range space) controls the smoothness of functions *f*_*k,p*_ and *GP*_*k*_, while the second penalty (null space) enables the removal of individual terms from the model entirely. The *γ* parameter can be used to globally increase the penalty and thus to obtain smoother, sparser and therefore potentially more robust models. Practical estimation was performed using techniques introduced in [58, 59] to increase computational speed and reduce memory requirements.

Six model specifications (Table 1) were considered for this analysis, varying by the definition of covariate effects in Equation 1 and whether the global GP term *GP*_*k,i*_(ℓ) was included. For each species, the final model (out of the six candidate models in Table 1) was selected based on its performance (AUC) on the test data (fold 5). Model 1 has no tuning parameters and was fit directly to the complete training data (folds 1-4). Models 2 through 6 were first tuned with respect to the global penalty *γ* ∈ {1, …, 4} based on 4-fold cross-validation on folds 1 through 4. Based on the value of *γ* that yielded the highest average AUC, the models were refit on the complete training data. All models were estimated using methods described in [60] to reduce run time and memory load.

**Table 1.**
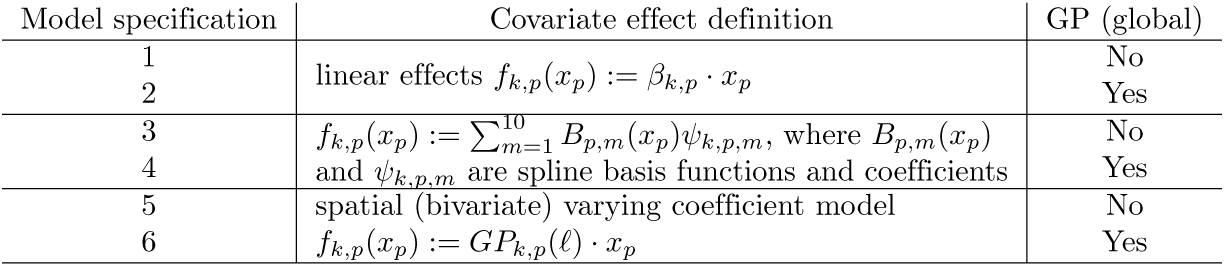
Model specifications considered in the analysis.

### Implementation

All calculations were performed using the **R** language environment [61]. Thematic maps were created using package **tmap** [62]. Data munging and pre-processing was performed using packages **dplyr** [63] and **tidyr** [64]. Spatial cross-validation was set up using package **blockCV** [65]. Package **mgcv** was used to fit the GAMs [56].

### Vectorial capacity of the mapped species

For each triatomine species that was mapped, information related to its importance in transmitting *T. cruzi* to humans was collated. The prevalence of infection with the *T. cruzi* parasite was calculated using the data from an existing repository [27]. Collections of less than twenty individuals of a species were excluded and the mean prevalence was calculated for all species where the remaining number of collections exceeded ten. Relevant behavioural data for each vector species was extracted from the literature.

## Results

### Species Distribution

A total of 30 species were mapped. A summary for all species that were modelled is provided in Table 2, including the specification of the model selected on training data as well as the AUC of this model evaluated on test data (fold 5) and the AUC obtained on the 20% randomly selected hold-out data (denoted by AUC*). The former is an indicator of the model’s transferability and ability to predict into new areas with potentially unseen covariate values or combinations, within the area that was modelled (cf. Figure 1). The latter value indicates how well the model interpolates as indicated by its ability to predict data points not used during model training and selection.

**Table 2.** Summary table for all species considered in the analysis ordered by number of presence observations.

| species | n presence<br>(test) | n background<br>(test) | Selected<br>model ($\gamma$) | AUC | AUC* |
| --- | --- | --- | --- | --- | --- |
| ***Triatoma infestans*** | 2499 (430) | 4458 (896) | 5 (3) | 0.96 | 0.97 |
| *Triatoma dimidiata* * | 1186 (154) | 2883 (787) | 6 (4) | 0.97 | 0.93 |
| ***Panstrongylus megistus*** | 968 (213) | 3409 (827) | 6 (1) | 0.83 | 0.89 |
| ***Triatoma brasiliensis*** | 820 (151) | 1989 (346) | 2 (1) | 0.69 | 0.81 |
| *Triatoma sordida* * | 809 (139) | 4919 (768) | 5 (2) | 0.83 | 0.89 |
| ***Triatoma pseudomaculata*** | 804 (239) | 3100 (687) | 4 (1) | 0.73 | 0.81 |
| ***Triatoma barberi*** | 455 (56) | 2161 (612) | 4 (1) | 0.88 | 0.92 |
| *Triatoma mexicana* | 430 (44) | 2046 (456) | 6 (2) | 0.92 | 0.96 |
| *Rhodnius prolixus* | 319 (42) | 962 (229) | 4 (2) | 0.64 | 0.85 |
| ***Panstrongylus geniculatus*** | 282 (39) | 6425 (824) | 3 (1) | 0.87 | 0.82 |
| *Triatoma longipennis* * | 282 (25) | 1974 (324) | 2 (2) | 0.97 | 0.95 |
| *Triatoma gerstaeckeri* | 280 (91) | 2480 (482) | 2 (1) | 0.85 | 0.96 |
| *Triatoma protracta* | 269 (42) | 2092 (212) | 6 (1) | 0.89 | 0.97 |
| *Triatoma pallidipennis* * | 256 (15) | 2286 (265) | 2 (2) | 0.96 | 0.96 |
| *Panstrongylus lutzi* | 136 (8) | 2965 (609) | 1 (–) | 0.78 | 0.92 |
| *Rhodnius pictipes* | 130 (25) | 3542 (940) | 5 (1) | 0.91 | 0.91 |
| *Triatoma rubida* | 126 (23) | 1913 (388) | 4 (1) | 0.98 | 0.95 |
| *Rhodnius neglectus* | 122 (18) | 3993 (655) | 4 (1) | 0.93 | 0.95 |
| *Triatoma mazzottii* * | 105 (7) | 2531 (581) | 4 (1) | 0.8 | – |
| *Rhodnius pallescens* | 93 (21) | 898 (202) | 2 (1) | 0.78 | 0.89 |
| *Triatoma guasayana* * | 92 (8) | 2761 (493) | 1 (–) | 0.52 | 0.75 |
| *Triatoma rubrovaria* | 92 (11) | 572 (227) | 3 (1) | 1.00 | 0.95 |
| *Rhodnius robustus* | 90 (22) | 3395 (915) | 4 (1) | 0.97 | 0.94 |
| *Triatoma sanguisuga* | 89 (17) | 701 (173) | 5 (2) | 1.00 | 0.92 |
| *Psammolestes tertius* | 75 (17) | 3969 (825) | 2 (3) | 0.79 | 0.90 |
| *Eratyrus mucronatus* * | 67 (13) | 2495 (656) | 5 (4) | 0.80 | 0.84 |
| *Triatoma maculata* | 65 (10) | 907 (169) | 5 (3) | 0.8 | 0.80 |
| *Panstrongylus chinai* | 57 (7) | 209 (53) | 4 (1) | 0.8 | – |
| *Rhodnius nasutus* | 56 (10) | 2178 (423) | 2 (1) | 0.90 | 0.83 |
| *Panstrongylus rufotuberculatus* * | 55 (13) | 3262 (608) | 3 (3) | 0.66 | 0.61 |

Maps for six example species, highlighting different genera, geographical ranges, dominant vector status and behaviours, are given in Figure 2. These are *Panstrongylus geniculatus, Panstrongylus megistus, Triatoma barberi, Triatoma brasiliensis, Triatoma infestans* and *Triatoma pseudomaculata*. Maps (including 95% CI) for all species listed in Table 2 are given in the supplement (S2 File, `.gri` file format). Respective visualisations are available from S3 File. The AUC values were all well above the 0.5 random classification threshold (mean: 0.85, SD: 0.12), indicating the maps usefulness to identify areas of higher probability of presence relative to background points. The comparatively low AUC values for species *T. brasiliensis* and *T. pseudomaculata* could be partially due to an overlap with many other species, thus making it difficult to discriminate between presence and background. The AUC* values were on average higher and had a lower variance (mean: 0.88, SD: 0.08) but generally consistent with the AUC values obtained on the test data (fold 5).

**Fig 2.**
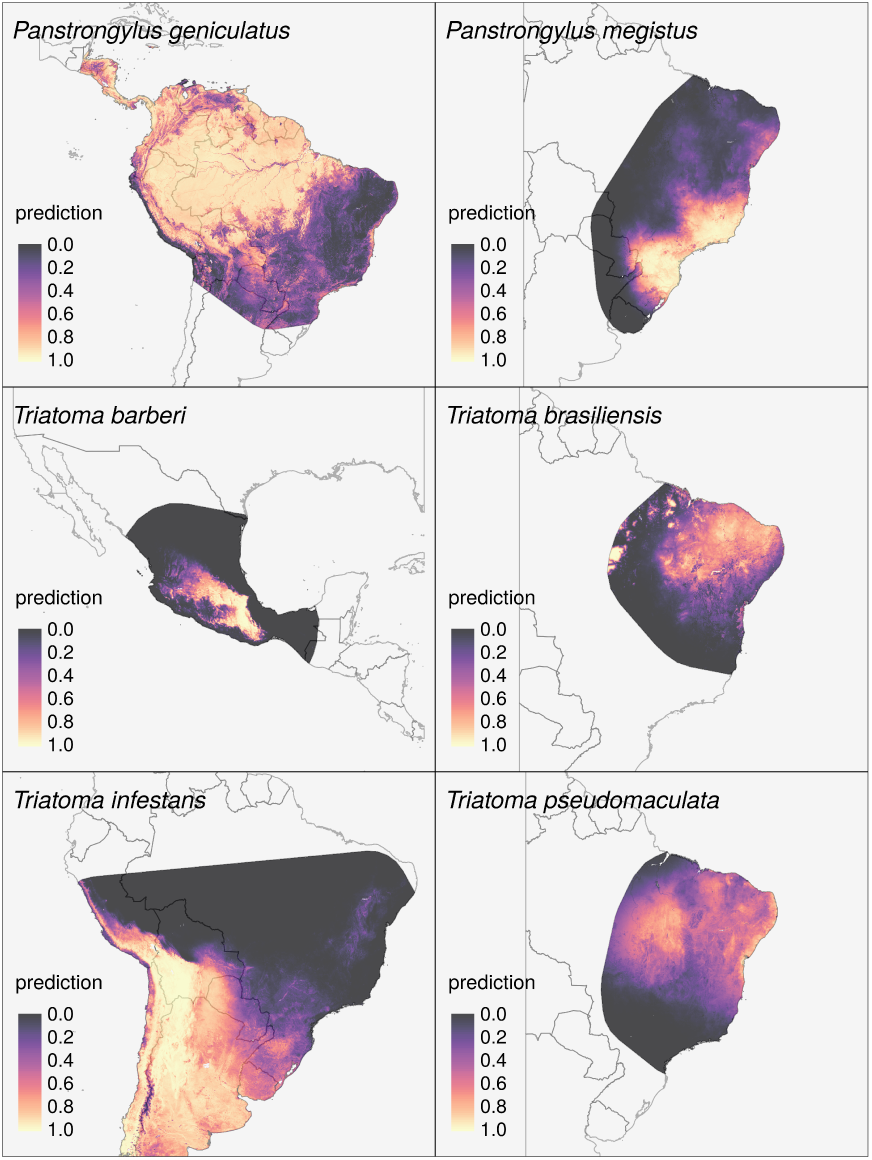
Predicted relative probability of occurrence at every 5 × 5 km pixel within the respective extended hull of species occurrence. The map for *T. brasiliensis* highlights some regions with potentially suitable habitats at the north and north-west of the extended hull, however, these regions were rarely sampled, containing neither presence nor background points and should be considered uncertain (cf. maps of lower and upper 95% confidence intervals in supplement S3 File). *T. infestans* presence locations on the other hand are mostly located close to the Andes with only a few background observations in the vicinity of these locations. The predictive map (Figure 2, bottom left panel) indicates a high probability of occurrence at high altitudes. This concurs with other studies that reported *T. infestans* presence in high altitudes [66, 67], however, difficult access means that the density of sampling points is often reduced in high altitude areas, which, in turn, exhibit higher uncertainty in the predictions.

### Vectorial capacity of the mapped species

The current state of knowledge on factors related to the capacity of each of the mapped species to transmit *T. cruzi* to humans (vectorial capacity) is summarised in Table 3. Specifically, mean infection prevalence, confirmation of human blood meals in natural vector populations, and confirmation of colonisation or invasion of homes (including in urban areas) are listed. Less information is available on the feeding-defecation interval or defecation location for each species, and these values may be influenced by the different experimental conditions used, so these variables are not included in Table 3 but sources of evidence are listed in File S4. The 30 most commonly reported species mapped here encompass the five most important dominant vectors that frequently colonise homes (*P. megistus, R. prolixus, T. brasiliensis, T. dimidiata* and *T. infestans*) as well as species that often colonise peridomestic habitats such as chicken coops, rats nests, boundary walls, wood piles, palm trees, and livestock housing. These species encompass a range of mean *T. cruzi* infection prevalences from 0.8 % in *T. sordida* to 55.6 % in *T. longipennis*, although for 13 of the most commonly reported species there was insufficient data to generate a reliable mean infection prevalence value. When viewing the summaries in Table 3, it is important to note that not all regions or species have been sampled or tested equally and a lack of published evidence for a specific component of vectorial capacity cannot be taken as definitive evidence of its absence. For example, no infections have been reported in *Eratyrus mucronatus* or *Psammolestes tertius*, but only 28 and 143 individuals have been tested, respectively, compared to 335,467 *T. sordida* individuals. In addition to the five important dominant vectors, there is evidence that many of the species mapped in this work are potential vectors of *T. cruzi*. Almost all of the 28 species that have been found to be infected with *T. cruzi* are known to invade homes, and at least 17 have been found to have fed on humans (Table 3). The sources of evidence — 126 published articles in total — are given in supplement File S4.

**Table 3.** Infection prevalence and behaviour of selected species.

| Species | Mean percent infected (n) | Human blood meals | Colonises homes | Invades home | Urban areas |
| --- | --- | --- | --- | --- | --- |
| *Eratyrus mucronatus* | NIF | - | yes | yes | - |
| *Panstrongylus chinai* | yes | - | yes | yes | - |
| *Panstrongylus geniculatus* | 7.6 (46) | yes | - | yes | yes |
| *Panstrongylus lutzi* | 6.8 (27) | yes | - | yes | - |
| *Panstrongylus megistus* | 8.3 (261) | yes | yes | yes | yes |
| *Panstrongylus rufotuberculatus* | yes | - | yes | yes | - |
| *Psammolestes tertius* | NIF | - | - | - | - |
| *Rhodnius nasutus* | 8.5 (73) | - | - | yes | - |
| *Rhodnius neglectus* | 4.0 (120) | yes | - | yes | yes |
| *Rhodnius pallescens* | yes | yes | - | yes | - |
| *Rhodnius prolixus* | 9.0 (10) | yes | yes | yes | - |
| *Rhodnius pictipes* | 24.6 (36) | yes | - | yes | yes |
| *Rhodnius robustus* | 25.9 (19) | - | - | yes | - |
| *Triatoma barberi* | yes | - | yes | yes | yes |
| *Triatoma brasiliensis* | 3.2 (918) | yes | yes | yes | - |
| *Triatoma dimidiata* | 28.9 (24) | yes | yes | yes | - |
| *Triatoma gerstaeckeri* | yes | yes | yes | yes | - |
| *Triatoma guasayana* | yes | yes | - | yes | - |
| *Triatoma infestans* | 27.6 (61) | yes | yes | yes | - |
| *Triatoma longipennis* | 55.6 (14) | - | yes | yes | yes |
| *Triatoma maculata* | 18.4 (18) | yes | - | yes | yes |
| *Triatoma mazzottii* | yes | - | - | yes | - |
| *Triatoma mexicana* | yes | - | - | yes | - |
| *Triatoma pallidipennis* | 4.8 (17) | yes | yes | yes | yes |
| *Triatoma protracta* | yes | - | yes | yes | yes |
| *Triatoma pseudomaculata* | 2.7 (988) | yes | yes | yes | yes |
| *Triatoma rubida* | yes | - | yes | yes | yes |
| *Triatoma rubrovaria* | 3.1 (17) | - | - | yes | yes |
| *Triatoma sanguisuga* | yes | yes | - | yes | yes |
| *Triatoma sordida* | 0.8 (1407) | yes | yes | yes | yes |

## Data availability

Original data sources for the construction of species occurrence and background data are publicly available from [26] and [27]. The predicted maps (rasters, i.e. the predicted values for every pixel) for all 30 species in Table 2 as well as the respective lower and upper bounds of the confidence intervals are available from [68] (S2 File). Visualisations of these data, i.e. map images, are available from supplement S3 File). The code for the analysis is available at https://github.com/adibender/chagas-vector-sdm. Since the occurrence data and covariate data cannot be accessed programmatically, this code provides full methodological details, but cannot be used to reproduce the entire analysis.

Summary table for thirty species for which predictive maps were created, ordered by number of occurrence observations. Species highlighted in bold are shown in Figure 2, maps of all 30 species (including 95% CI) are given in the supplement (S2 File and S3 File). Asterisks (*) mark species for which the block width *w*_*k*_ used to construct spatial blocking was smaller than a preliminary estimation of the range of spatial auto-correlation, thus estimated AUC values might be overoptimistic in these cases. Column “Selected model” indicates the model specification that was selected based on its performance on the training data and refers to the model specifications defined in Table 1. If applicable, *γ* indicates the value of the selected global penalty multiplier (cf. Eq. (2)). The reported AUC value was calculated on the test data set (fold 5 in Figure 1). The AUC* value was calculated on the 20% hold-out data (not spatially blocked). Entries “–” indicate that there were not enough observations and/or unique predicted values in the hold-out data for calculation.

The mean infection prevalence is given together with the number of triatomine collections that contributed to the mean. If there were insufficient data to calculate the mean, i.e. fewer than 10 collections of ≥ 20 individuals, then records of infected individuals (yes) or no infections found (NIF) are noted. Instances where there was no evidence for a particular behaviour are denoted by “-”.

## Discussion

This study models the contemporary geospatial distributions of the thirty most commonly reported triatomine species and putative vectors of the *T. cruzi* parasite to humans. Our approach allows the distributions of these different species to be compared, and to be overlaid, which increases our understanding of the community of vector species at different locations in the current intervention era.

Our aim was to consider the most commonly reported species in Latin America. To provide policy makers, stakeholders and researchers with relevant information, we included all species for which distribution maps could be reasonably estimated. However, as can be seen from Table 2, the training (and test) data for many species contained fewer than 300 or even fewer than than 150 presence observations reported since the year 2000. For some of these species, the test fold (cf. Table 2) only contained a few presence locations, sometimes concentrated in a few blocks, i.e., within a small geographical area. AUC values will thus tend to be less robust and potentially over- or under optimistic as sample size decreases. One should therefore be particularly cautious when interpreting results of species with few presence data points and always take into account the uncertainty of the estimates as presented in the supplement (S3 File). There are locally important species for which maps could not be produced because they are only found within areas where the relevant surveillance records are not publicly available or because their range is limited so only small numbers of observations exist. For example, *Rhodnius ecuadoriensis* is an important vector in Ecuador [69] but the databases used in this study only provided 11 and 23 records, respectively, for known collection dates after the year 2000.

Earlier studies [19, 20, 22, 23, 25], most notably [24], have modelled the distributions of some of these species but only in specific regions, states or countries. Additionally, comparisons with the previous work are limited because of differences in methodology, data sets and spatial extent under consideration. Only visual comparison is possible in most cases because the predicted values generated by other studies are not openly available, precluding quantitative assessment of the different versions. In general, our respective predicted maps show very good agreement with respect to regions that highlight higher vs. lower probabilities of occurrence, but often differ with respect to the spatial extent of the region modelled.

The maps generated by this study should be considered in the context of the behaviour and vectorial capacity of each species. Summaries of the available evidence are provided here and show that most of these species are potentially important vectors of *T. cruzi* to humans. Each of the indicators of vectorial capacity summarised at a species level by this study may vary within the range of the species, as well as between species [70, 71]. It is therefore important to map spatial variation in these characteristics, as well as in the species themselves, in order to identify where regions of vectorial transmission risk are likely to exist.

## Supporting information

### S1 File. Presence/background data and spatial blocking

This file contains one figure for each species contained in Table 2. In each figure, the left panel shows raw presence absence data constructed as described in Figure 1, panels (A) through (C). Panel (D) shows the spatial allocation of blocks that define training (folds 1-4) and test (fold 5) data. The figures are published in [72] and available from https://figshare.com/s/f027db53093230373fa5.

### S2 File Predicted 5 × 5 km resolution maps (raw)

Files in `.gri` format that contain the predicted relative probability of occurrence for 30 traitomine species (Chagas vectors). The files consist of the predicted probabilities (`predicted-maps-tgb.gri`) and the respective lower (`predicted-cil-tgb.gri`) and upper (`predcited-ciu-tgb.gri`) confidence intervals. Each layer in the raster brick represents one species. The files are published in [68] and available from https://figshare.com/s/78e2c83427772ddd6cc9.

The data can be read into **R** using command `raster::brick`. For example:

~~~
# read in predcitions for all 30 species
pred_maps <-raster::brick(“predicted-maps-tgb.grd”)
# visualise prediction for species Panstrongylus megistus
sp::plot(pred_maps[[“Panstrongylus.megistus”]])
~~~

### S3 File. Predicted 5 × 5 km resolution maps (visualisation)

This file contains one figure for each species contained in Table 2 depicting the predicted relative probabilities of occurrence for each species. In each figure, the left and right panel show the lower and upper bound of the 95% confidence interval, respectively. The middle panel the expected value. The file is published in [73] and available from https://figshare.com/s/ffeeb36f30dc6c128819.

### S4 File. Sources of evidence for variables linked to vectorial capacity

This file contains the full information summarised in Table 3 together with citations for the sources of evidence used. The file is available from https://figshare.com/s/3dd02aa5a969aefb08ad.

## Acknowledgements

This work was funded by a Bill & Melinda Gates Foundation grant (OPP1053338) to S.W.L. and a Wellcome grant (108440/Z/15/Z) to C.L.M.

